# Venular-centered thrombo-inflammation drives microvascular failure after arterial recanalization in acute mesenteric ischemia: a translational study

**DOI:** 10.64898/2026.01.08.698481

**Authors:** Déborah François, Lambert Kernanet, Abigail Brami, Véronique Arocas, Marie-Christine Bouton, Dominique Cazals-Hatem, Kevin Guedj, Benoit Ho Tin Noe, Olivier Corcos, Yacine Boulaftali, Alexandre Nuzzo

## Abstract

Acute mesenteric ischemia (AMI) remains associated with high mortality despite prompt revascularization, suggesting that downstream ischemia–reperfusion injury contributes to poor outcomes. However, the microvascular mechanisms underlying this process remain poorly defined.

We analyzed admission blood samples from patients with arterial AMI and non-ischemic controls and investigated thrombo-inflammatory responses in a murine superior mesenteric artery occlusion (SMAO) model. In mice, intravital microscopy was used to directly visualize mesenteric microcirculatory flow and thrombo-inflammatory events during ischemia–reperfusion.

Patients with AMI displayed a marked systemic thrombo-inflammatory profile, characterized by elevated inflammatory markers, neutrophil activation, platelet activation, and alterations in coagulation-related proteins, which were closely mirrored in the SMAO model. Intravital microscopy revealed a dissociation between arterial and microvascular reperfusion: while arteriolar flow partially recovered after recanalization, venular perfusion remained severely impaired and was associated with early blood cell stasis and stable venular thrombus formation. Thrombi developed through a sequential process initiated during ischemia and amplified during reperfusion.

Together, these findings identify venular-centered thrombo-inflammation as a key determinant of microvascular dysfunction and intestinal injury in arterial AMI, and provide a mechanistic framework for targeting thrombo-inflammatory pathways beyond arterial reperfusion that may extend to other clinical forms of AMI, including non-occlusive mesenteric ischemia (NOMI).

## INTRODUCTION

Acute mesenteric ischemia (AMI) is a life-threatening gastrointestinal emergency that remains associated with a mortality rate of 30-50%, despite advances in diagnosis, arterial revascularization techniques, and the development of specialized referral centers^1–3^. Prompt restoration of arterial blood flow is essential to prevent transmural intestinal necrosis and its systemic complications, including sepsis and multiorgan failure. However, clinical outcomes remain poor even after technically successful revascularization, indicating that downstream mechanisms beyond large-vessel recanalization critically contribute to intestinal injury and prognosis.

Ischemia–reperfusion (IR) injury is increasingly recognized as a central driver of tissue damage and remote multiorgan failure in AMI. Although reperfusion is required to salvage ischemic bowel, the abrupt restoration of blood flow paradoxically exacerbates injury through oxidative stress, endothelial dysfunction, and immune activation. In the intestine, IR compromises epithelial integrity and barrier function, facilitating bacterial translocation and triggering systemic inflammatory responses that may rapidly progress to organ failure^4^. Despite its clinical relevance, no targeted therapies currently exist to mitigate IR in AMI. Management remains largely focused on restoring vascular patency, with no interventions aimed at enhancing intestinal tolerance to ischemia or promoting recovery after reperfusion.

Experimental models of superior mesenteric artery occlusion (SMAO) have shown that inflammation is an early and sustained feature of both ischemia and reperfusion.^5,6^ Hypoxia-induced oxidative stress during ischemia is amplified upon reperfusion by oxygen influx and activation of endothelial cells, neutrophils, macrophages, and platelets, resulting in profound microvascular dysfunction^7^. Growing evidence indicates that inflammatory and thrombotic pathways are tightly interconnected in this setting, forming a thrombo-inflammatory response that amplifies tissue injury. Endothelial cell-mediated T-cell–dependent neutrophil recruitment^8^, excessive ROS production^9^, neutrophil extracellular traps (NETs) formation^10,11^, and platelet–fibrin deposition^12–16^, have all been implicated in intestinal IR injury, yet these processes are often studied separately. A similar thrombo-inflammatory cascade has been well described in ischemic stroke, where successful large-artery recanalization does not necessarily restore effective microvascular perfusion or clinical recovery. In animal models of transient middle cerebral artery occlusion, intravital microscopy has shown that thrombo-inflammation is initiated during ischemia and exacerbated upon reperfusion, leading to persistent microvascular obstruction dominated by leukocyte–platelet aggregates, fibrin deposition, and stable venular thrombi^17,18^. These insights have reshaped the understanding of post-ischemic tissue injury and highlighted microvascular thrombo-inflammation as a therapeutic target. Given the intestine’s unique exposure to luminal microbes and its close integration with the immune system, thrombo-inflammation is likely to play an even more deleterious role in AMI-associated IR yet remains poorly characterized.

In this study, we combined a translational analysis of thrombo-inflammatory plasma biomarkers in patients with AMI from the SURVIBIO biobank with mechanistic investigations in a standardized murine model of superior mesenteric artery occlusion. Using intravital microscopy, we examined the dynamics of intestinal microvascular thrombo-inflammation during IR. Together, these complementary human and experimental approaches were designed to improve the pathophysiological understanding of AMI and to explore thrombo-inflammatory pathways as potential therapeutic targets beyond arterial recanalization.

## METHODS

### Human plasma collection

Patients were prospectively enrolled in the SURVIBIO diagnostic study (NCT03518099), a cross-sectional study including patients with AMI and controls presenting with non-ischemic acute abdominal pain, as previously described^19^. Patients with AMI were managed at the intestinal stroke center (SURVI, Beaujon Hospital, Clichy, France), a tertiary referral center providing 24/7 standardized multidisciplinary care^20^. Standard treatment included, aspirin, heparin, antibiotics, and urgent endovascular or surgical revascularization^20,21^. In patients with suspected transmural necrosis, laparotomy was performed to assess bowel viability and guide intestinal resection^22^. Serum samples from 20 patients (10 with arterial AMI and 10 non-AMI controls) included between January 2016 and March 2018 were selected for analysis. Baseline demographic, clinical, and biological data were collected at admission. Blood samples were obtained on admission, immediately centrifuged, and serum aliquots were stored at-80°C for biomarker analyses. Within an untargeted proteomics discovery workflow, we specifically examined thrombo-inflammatory markers including P-selectin and myeloperoxidase (MPO). Detailed proteomics methods are provided in Supplementary Data 1. The study was approved by the Ethics Committee of Paris-Nord Val de Seine University Hospitals, and all participants provided written informed consent prior to enrollment.

### Superior Mesenteric Artery Occlusion (SMAO) mouse model

Before surgery, all mice received a s.c. injection of analgesia (1mg/kg buprenorphine) at least 30 minutes before anesthesia. After the isoflurane induction at 4%, anesthesia maintenance was achieved by 1.5% isoflurane via facemask during SMAO (2 hours).

Temperature homeostasis was achieved through the use of a thermostated heating plate. Using aseptic technique, a midline laparotomy was performed, the intestines were eviscerated and the mesentery exposed. The superior mesenteric artery was identified and clamped using an atraumatic microvascular clamp. Following 60 minutes of intestinal occlusion, the atraumatic clamp was removed. During the occlusion (1hour) and recanalization (1hour) phases, the intestine and mesentery were constantly hydrated with sterile NaCl 0,9%. Following surgery, animals were sacrificed by cervical dislocation. A group size of 8 to 15 animals per group, sham (laparotomy without vascular occlusion) or SMAO mice, was determined to optimize the animal model taking into account the statistical risk of error (5%) and the power of the tests (80%).

All procedures adhered to the ethical guidelines of the European Directive (2010/63/EU) and were approved by the ethical committee of the French Ministry of Higher Education and Research. The experiments followed the Animal Research: Reporting *In Vivo* Experiments 2.0 guidelines and were approved under the reference number n°40873 & 46125. All animals were maintained under standard conditions and kept on a 12-hour light/dark cycle with food and water given ad libitum (maximum 5 animals per cage). C57BL/6J female mice 8 to 12 weeks old provided by Janvier laboratory, were allowed an acclimatization period for at least 1 week before the experiment.

### Real-Time Intravital Imaging

Mesenteric microcirculation on distal arterioles (50-100µm) and venules (100-250µm) couples of the superior mesenteric artery was visualized using a fluorescence macroscope (Z16 APO, Leica Microsystems) equipped with a thermostated heating plate and an objective connected to a CMOS camera (ORCA-Flash 4.0 LTPlus; Hamamatsu photonics). Data acquisition and analysis were done using Micromanager (Open Source Software). All fluorescent markers were administrated intravenously into the retroorbital sinus. Rhodamine 6G was used to label platelets and leukocytes, AlexaFluor488 (Invitrogen)-conjugated rat anti-mouse Ly6G (BioXCell) was used to stain neutrophils, AlexaFluor555 (Invitrogen)- conjugated rat anti-mouse GPIX (Emfret) was used to stain platelets and AlexaFluor647 (Invitrogen)-conjugated mouse anti-mouse anti-fibrin (Sigma) was used to stain fibrin. To monitor cell recruitment kinetics during ischemia and recanalization, 10 seconds intravital imaging acquisitions were performed before SMAO, at T0, T20, T40 and T60 minutes after SMAO and at T0, T20, T40 and T60 minutes after recanalization. For all timepoints, leukocytes number per field of view were counted in the arterioles (10-second acquisition time) and a global fluorescence was measured in the venules (3 measures per acquisition). Venous thrombosis occurrence was assessed in each group and is expressed as percentage of mice developing venous thrombosis during recanalization after SMAO.

### Laser Doppler Interferometry

Intestinal mesenteric perfusion was analyzed through blood cell velocity in micro-vessels measured using a single-point laser Doppler vibrometer with an integrated CCD video camera (Polytec). To allow specific measurement of blood cell velocity, breathing-related and flow-induced vessel wall vibrations were identified by their bidirectional associated signal and eliminated by applying a low-pass filter set at 2 kHz. Frequencies due to the unidirectional out-of-plane vibrations caused by circulating blood cells were converted to speed according to the formula v=(Δfxλ)/(2xcosα), where v is the blood cell velocity, Δf is the Doppler-frequency shift, λ is the wavelength of the emitted wave (633 nm), and α is the angle between the blood cell direction and the incident laser beam, which was estimated at 80°. Data were recorded and treated using the Polytec Vibrometer Software. Perfusion images were acquired at baseline (before SMAO), at T0, T20, T40 and T60 minutes after SMAO and at T0, T20, T40 and T60 minutes after recanalization. Perfusion data was expressed as a percentage of baseline.

### Mice histological staining, scoring and intestinal damages quantification

Following euthanasia of mice groups, segments of intestine were harvested, fixed in 4% paraformaldehyde and embedded in Paraffin for histological analysis. Intestinal sections of 5µm thickness were stained with hematoxylin and eosin and subjected to histological scoring to evaluate tissue damage. Mucosal damage score was calculated as described by Chiu *et al* ^23^. Macroscopic assessment of intestinal damaged sections was performed by measuring damaged sections along the entire length of the extracted intestine and is expressed as a percentage of total intestinal length. Chiu score and quantification of intestinal damages was assessed blindly by two pathologists.

### Immunofluorescence staining

For immunofluorescent staining, paraformaldehyde fixed paraffin sections of intestine and thrombotic mesenteric vessels were subjected to rehydration and antigen retrieval. After blocking with PBS/BSA 3% for an hour, the sections were incubated overnight with primary antibodies. The following day, sections were washed and incubated with secondary antibodies for 2h. Stained sections were mounted using Fluoromount containing DAPI (Abcam). The following antibodies were used: Rabbit anti-mouse CD42b (Abcam), Rat anti-mouse CD45 (Biolegend), Rabbit anti-mouse Fibrin/Fibrinogen (Dako), Rat anti-mouse LY6G (BD pharmingen), Mouse anti-mouse occludin (Invitrogen) and Goat anti-LPS (Invitrogen). Secondary antibodies were Goat anti-rabbit Cy3, Goat anti-rat 488, Goat anti-mouse 488 (Jackson Immuno Research).

### Enzyme-linked immunosorbent assay (ELISA)

Plasma samples from sham or SMAO mice group were prepared and used to assess levels of InterLeukine-6 (IL-6) (Invitrogen), Myeloperoxydase (MPO - R&Dsystem), soluble P-selectine (sP-sel - R&Dsystem) and TAT complex (AssayMax) according to the manufacturer’s specifications.

### Statistical analysis

Data are presented as means ± SEM. SMAO mice data were compared to Sham mice data using a nonparametric Mann-Whitney statistic test. Values of *p*<0.05 were considered statistically significant. For patient’s characteristics, continuous variables were presented as median (IQR). Variables were compared using the Mann–Whiney test. All tests were two-sided and p ≤0.05 was considered significant.

## RESULTS

### Thrombo-inflammation biomarkers in patients and mice

We analyzed admission blood samples from 20 patients enrolled in the SURVIBIO biobank (NCT03518099), including 10 patients with confirmed arterial AMI and 10 controls admitted for non-ischemic acute abdominal pain (Table 1). The final diagnoses among the 10 control patients were mechanical small bowel obstruction (n=6), irritable bowel syndrome (n=3), and renal colic (n=1). Among AMI patients, the median age was 69 years (IQR 60–85), 50% were female, and arterial occlusion was attributed to atherosclerosis in 80% and embolism in 20%. Compared with controls, AMI patients exhibited a marked systemic inflammatory response at presentation, with significantly higher circulating levels of C-reactive protein (*p*<0.001), serum amyloid A1 (*p*=0.001), and procalcitonin (*p*=0.002) (Table1 & Figure 1). Leukocyte counts were increased (*p*=0.01), with trends toward higher neutrophil counts (*p*=0.02) and MPO levels (*p*=0.05). Markers of platelet activation and thrombo-inflammation were also elevated, including P-selectin (*p*=0.02) and plasminogen activator inhibitor-1 (PAI-1; *p*=0.14). In parallel, fibrinogen γ chain levels were significantly increased (*p*=0.001), whereas fibrinogen β chain levels and platelet counts did not differ significantly. Fibrinogen α chain, prothrombin, factor XIII, and citrulline levels tended to be lower in AMI patients (Table 1 & Figure 1).

**Figure 1:**
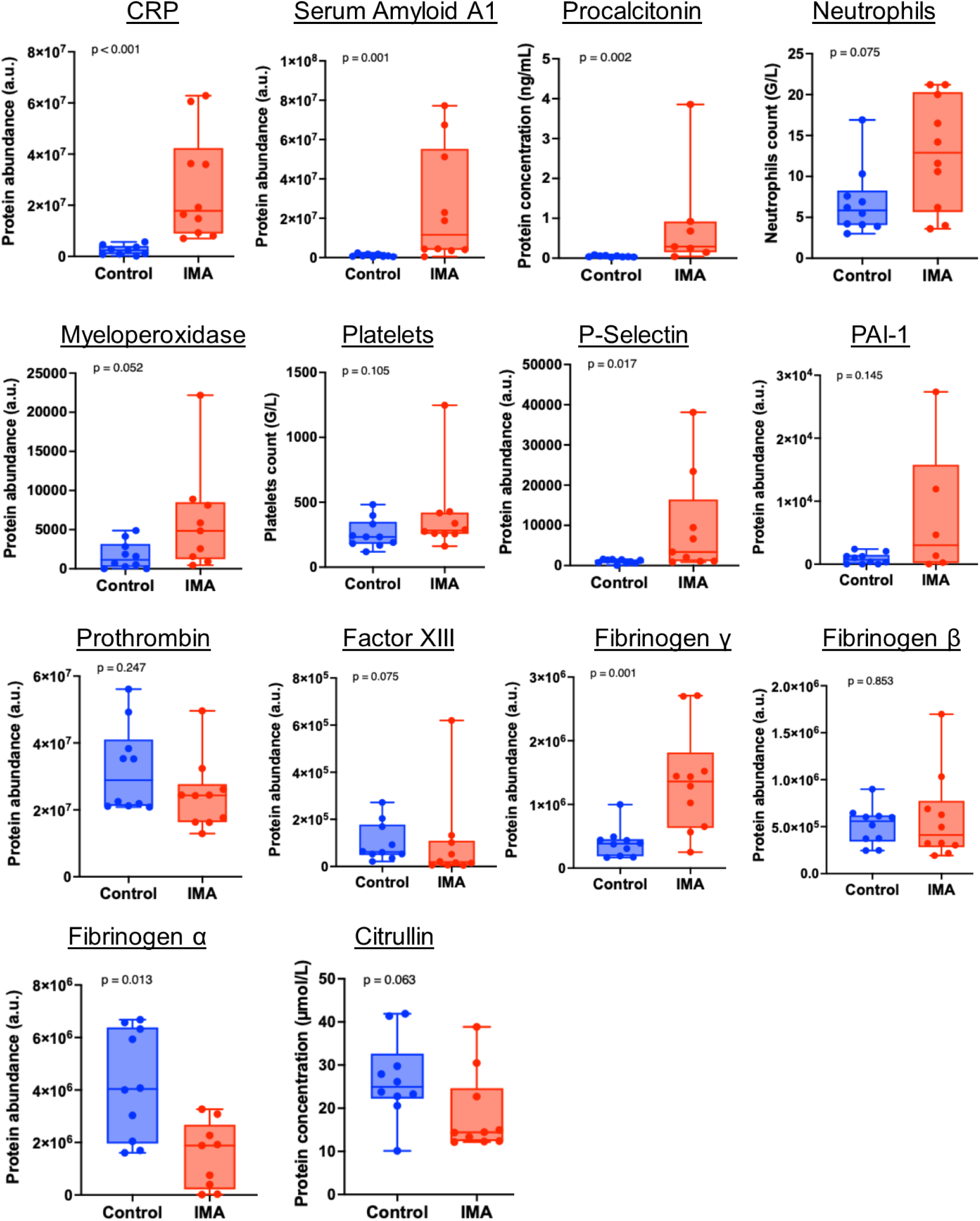
Biomarkers of thrombo-inflammation and epithelial damage in AMI patients. Biomarkers of thrombo-inflammation and epithelial damage in AMI patients. Proteomics data are expressed in arbitrary units (a.u). Quantitative variables are expressed as median and IQR. *p*≤0.05 was considered significant. *CRP: C-reactive protein*

**Table 1:** Characteristics of the patient’s cohort Abbreviations: IQR: interquartile range; SOFA: Sequential Organ Failure Assessment.

|  | Control<br>patients<br>n=10 (%) | AMI<br>patients<br>n=10 (%) | *p-value*<sup>1</sup> |
| --- | --- | --- | --- |
| Age, years, median (IQR) | 37 (27-72) | 69 (60-85) | 0.01 |
| Female | 3 (30) | 5 (50) | 0.65 |
| Atherosclerosis (vs. arterial embolism) | - | 8 (80) | - |
| *Comorbidities* |  |  |  |
| Tobacco use | 3 (30) | 3 (30) | 1.00 |
| Arterial hypertension | 2 (20) | 6 (60) | 0.17 |
| Dyslipidemia | 0 (0) | 4 (40) | 0.09 |
| Diabetes mellitus | 1 (10) | 4 (40) | 0.30 |
| Myocardial ischemia | 0 (0) | 2 (20) | 0.47 |
| Stroke | 0 (0) | 2 (20) | 0.47 |
| Limb ischemia | 0 (0) | 2 (20) | 0.47 |
| Atrial fibrillation | 1 (10) | 3 (30) | 0.58 |
| *Clinical and biological presentation* |  |  |  |
| Guarding | 3 (30) | 2 (20) | 1.00 |
| Organ failure (SOFA>2) | 3 (30) | 5 (50) | 0.37 |
| Leukocytes (G/L) | 8.8 (7.4-12.5) | (10.5-19.1) | 0.01 |
| Neutrophils (G/L) | 6.9 (4.5-10.1) | 10.8 (8.6-16.6) | 0.02 |
| Lymphocyte (G/L) | 0.9 (0.7-1.0) | 1.2 (0.8-1.6) | 0.35 |
| Platelets (G/L) | 232 (191-395) | 307 (217-418) | 0.42 |
| C-reactive protein (mg/L) | 4.5 (1.0-21.5) | 106.0 (26-192.5) | <0.001 |
| Lactate (mmol/L) | 1.6 (1.3-2.6) | 1.5 (0.9-2.6) | 0.64 |
| Creatinine levels (μmol/L) | 72 (63-103) | 70 (60-86) | 0.32 |
| *Follow-up* |  |  |  |
| ICU admission | 0 (0) | 5 (50) | 0.03 |
| Abdominal surgery | 2 (20) | 4 (40) | 0.63 |
| 12-month mortality | 0 (0) | 2 (20) | 0.47 |

To assess these thrombo-inflammatory alterations under standardized experimental conditions, blood samples were collected in mice subjected to superior mesenteric artery occlusion (SMAO; n=9) or sham surgery (n=7). Complete blood counts were comparable between groups, except for a significant reduction in platelet counts (*p*=0.02) in SMAO mice. Consistent with the human data, circulating P-selectin (*p*=0.02) and MPO (*p*=0.04) levels were increased. In addition, IL-6 levels were markedly elevated (*p*<0.001), and thrombin–antithrombin (TAT) complexes were increased during IR (p = 0.04) (Figure 2).

**Figure 2:**
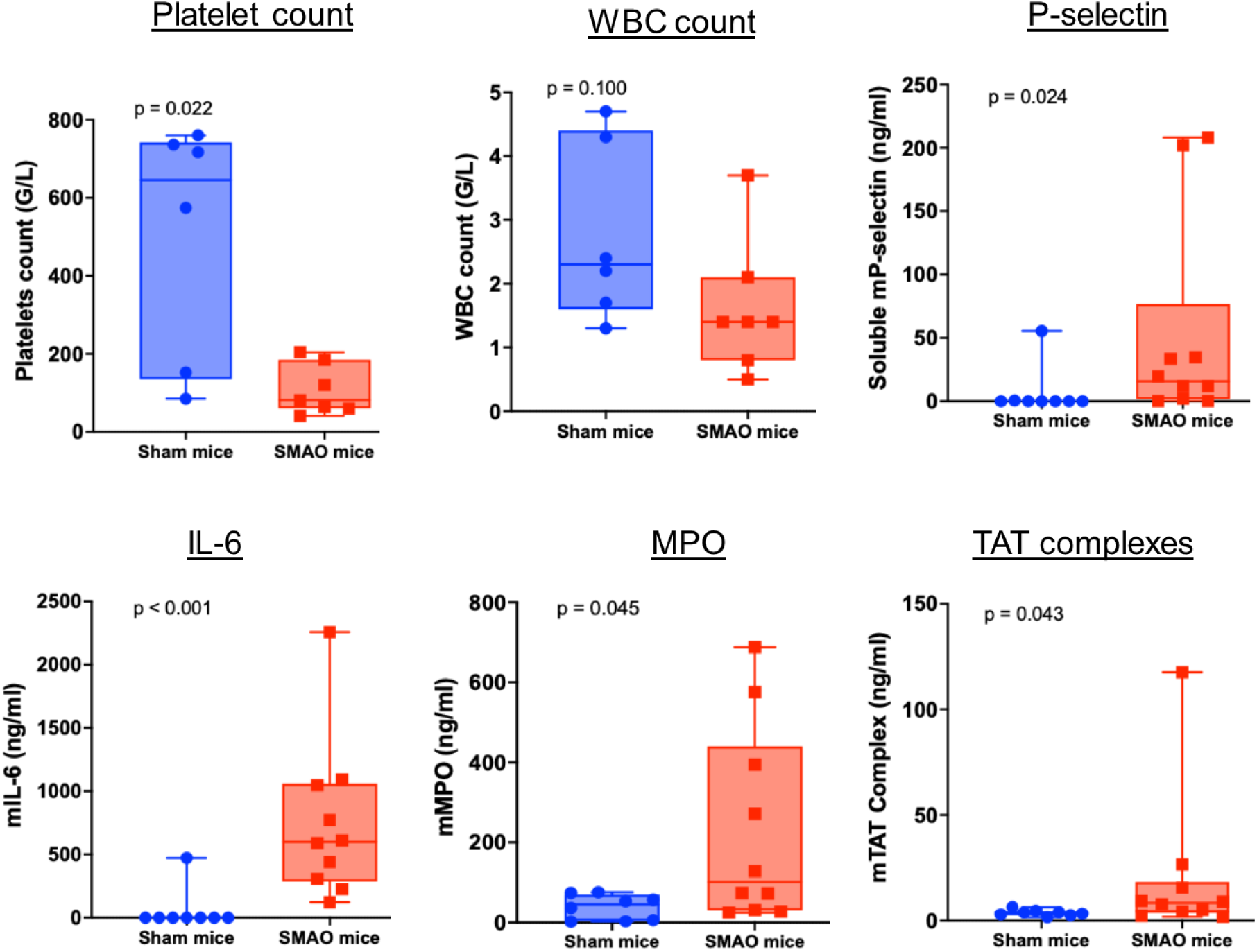
Biomarkers of thrombo-inflammation in mice. Characteristics and inflammation and thrombosis markers of control or SMAO mice. Quantitative variables of platelets and white blood cells (WBC) counts are expressed as median and IQR (n=6 Sham mice and n=7 SMAO mice). Variables were compared using the Mann–Whiney test. All tests are two-sided. Plasmatic markers (IL-6, Myeloperoxydase (MPO), Soluble P-selectin and TAT Complexes) are expressed as mean ± SEM (n=8 Sham mice and n=14 SMAO mice). *p*≤0.05 was considered significant.

### SMAO disrupts downstream microvascular flow and leads to venous thrombosis after recanalization

To investigate the microvascular origin of thrombo-inflammatory signatures observed in human AMI and in the SMAO model, we used intravital microscopy to visualize mesenteric microcirculatory flow and thrombo-inflammatory events during IR.

Following superior mesenteric artery occlusion, arteriolar blood flow, assessed by red blood cell velocity, decreased to 27 ± 4% of baseline and partially recovered to 72 ± 19% after 20 minutes of reperfusion (Figure 3A). However, arteriolar blood flow progressively declined thereafter, reaching 47 ± 12% at 60 minutes of reperfusion. Leukocyte rolling and adhesion along arteriolar walls increased during ischemia and progressively resolved during reperfusion, becoming undetectable after 40 minutes (Figure 4A & 4C). In contrast, venular blood flow decreased to 41 ± 6% of baseline during ischemia but remained severely impaired throughout reperfusion (46 ± 6% at 60 minutes; Figure 3B). Reperfusion was associated with a rapid and sustained increase in venular blood cell stasis, reflected by a 3.6-fold rise in global fluorescence intensity (*p*=0.007) after 60 minutes (Figure 4B-C), and with the formation of venular thrombi in 77% of SMAO mice (Figure 4D). Thrombi appeared as early as 20 minutes after reperfusion, progressively enlarged, and remained stable throughout the observation period. No arterial thrombi were detected. In the remaining mice, persistent platelet and leukocyte adhesion was observed without overt thrombus formation. No microvascular alterations were observed in sham mice (Figure 4).

**Figure 3:**
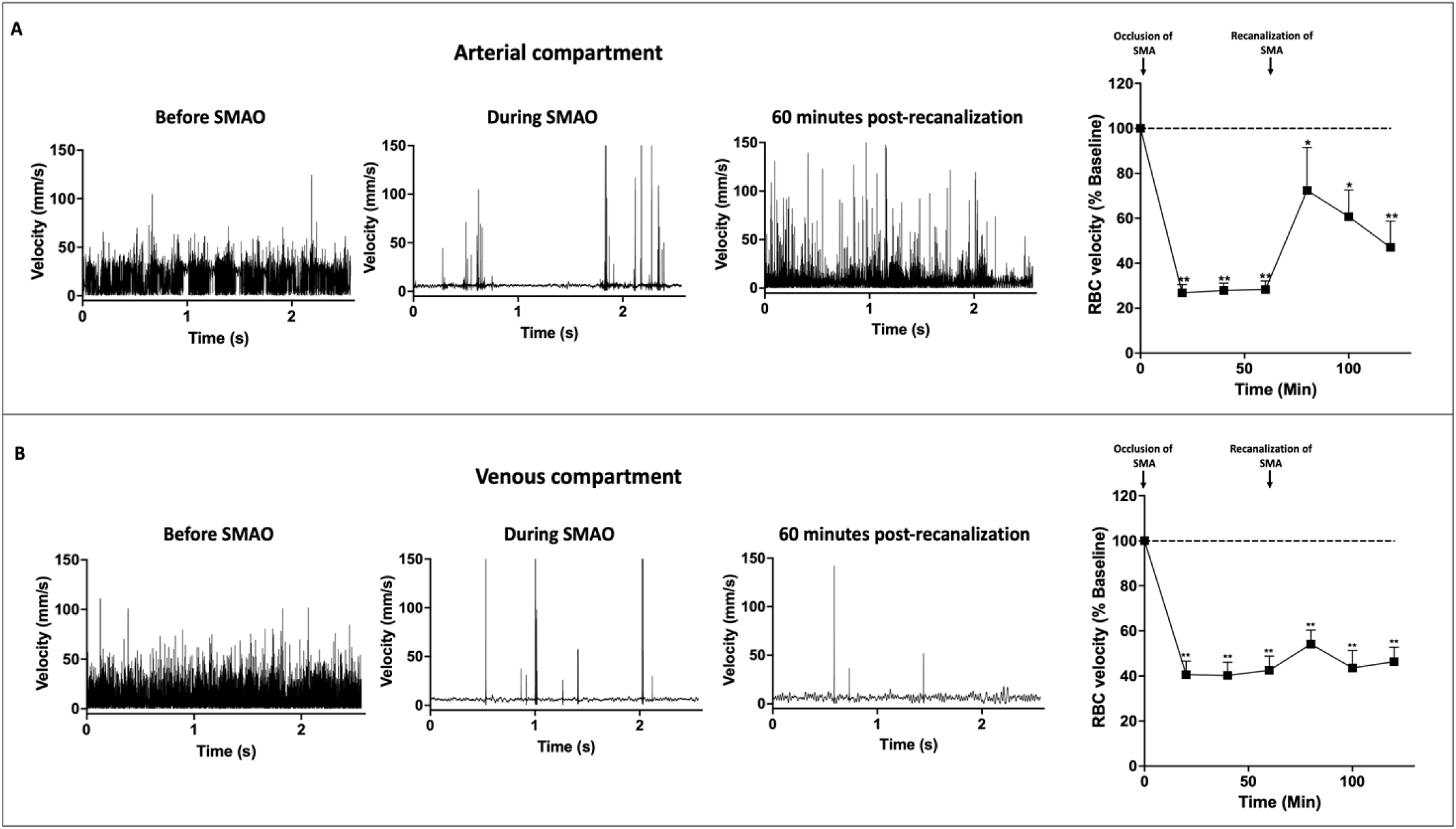
Microvascular blood flow during intestinal ischemia-reperfusion. **(A)** Arterioles and **(B**) venules variation in red blood cell velocity at baseline, during superior mesenteric artery (SMA) occlusion (0, 20, 40 & 60 minutes after SMA occlusion) and after SMA recanalization (20, 40 & 60 minutes). Results are expressed as percentages relative to baseline values. n=7 mice on, 7 arterioles (50 to 100µm diameter) and 7 venules (150 to 300µm diameter). Error bars represent SEM. \**p*<0.05 and \*\**p*<0.01 compared with baseline.

**Figure 4:**
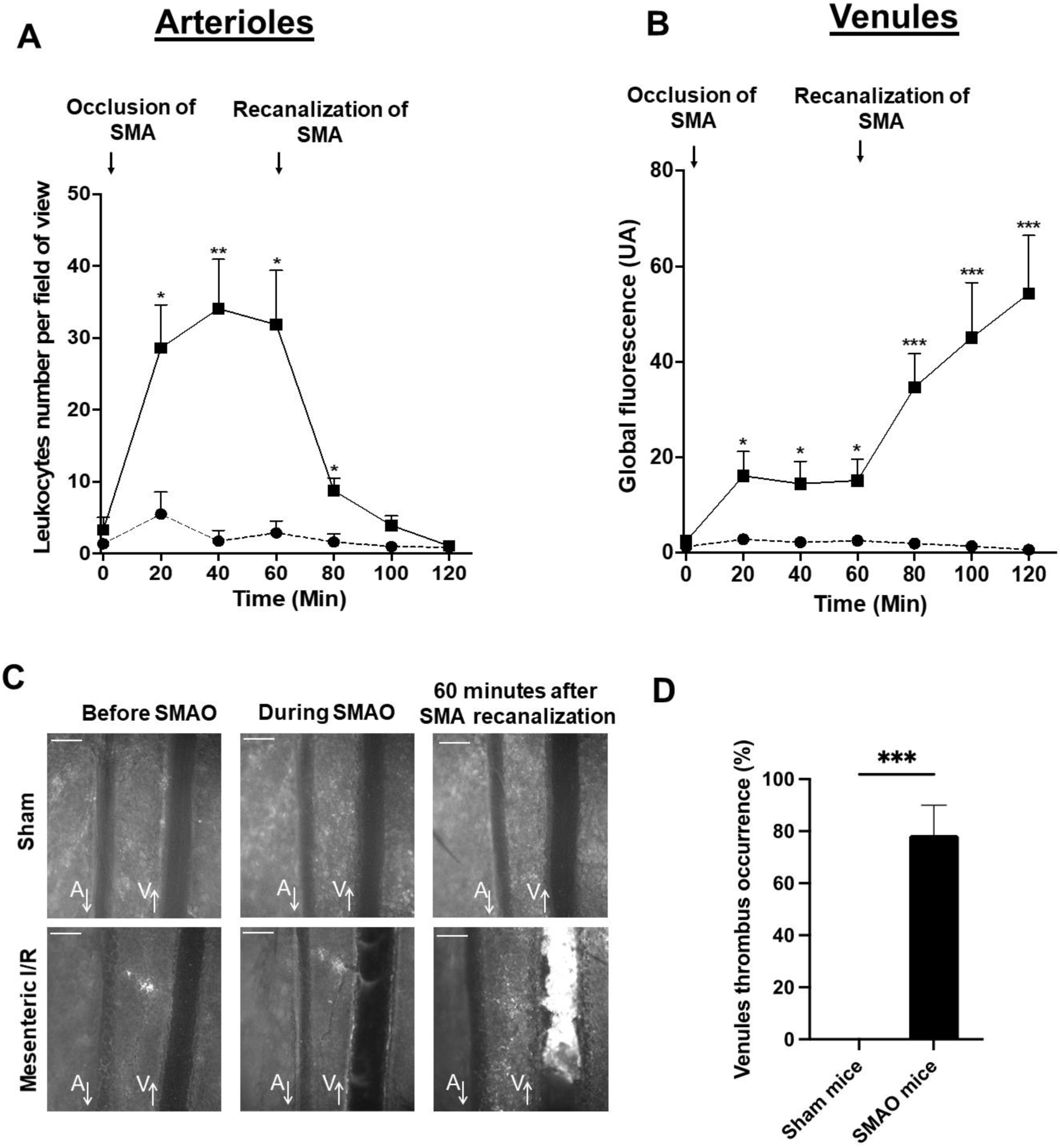
Microvascular alterations following superior mesenteric artery occlusion. **(A)** Quantification of leukocyte adhesion in mesenteric arterioles before ischemia, during superior mesenteric artery occlusion (SMAO), and after arterial recanalization. Leukocytes were counted per field of view over a 10-s intravital microscopy recording. **(B)** Quantification of blood cell stasis in mesenteric venules, expressed as global fluorescence intensity normalized to venular area, measured before ischemia, during SMAO, and after recanalization (three measurements per 10-s recording). **(C)** Representative fluorescent intravital microscopy images showing rhodamine 6G–labeled platelets and leukocytes adhering to arteriolar and venular walls downstream of the superior mesenteric artery. Images were acquired before SMAO, immediately before recanalization, and 60 min after recanalization. Arrows indicate blood flow direction. Scale bar, 200 μm. **(D)** Incidence of venular thrombus formation 60 min after reperfusion, assessed by intravital microscopy. Arterioles analyzed measured 50–100 μm in diameter, and venules 150–300 μm. Data are shown as mean ± SEM. Sham, n = 9; SMAO, n = 14. \**p*< 0.05, \*\**p*< 0.01, \*\*\**p*< 0.001 versus baseline.

Fluorescent intravital microscopy and analysis of venular thrombus sections further characterized the cellular dynamics of thrombus formation. Platelet stagnation occurred early after arterial occlusion, whereas neutrophils initially localized predominantly to the perivascular space (Figure 5A). During ischemia, platelet clusters progressively formed and colocalized first with fibrin and subsequently with neutrophils. These aggregates expanded during reperfusion, giving rise to stable venular thrombi. Immunofluorescence analysis revealed a thrombus architecture composed of a platelet-and fibrin-rich core with entrapped red blood cells, surrounded by an outer shell enriched in fibrin, neutrophils, and platelets (Figure 5B).

**Figure 5:**
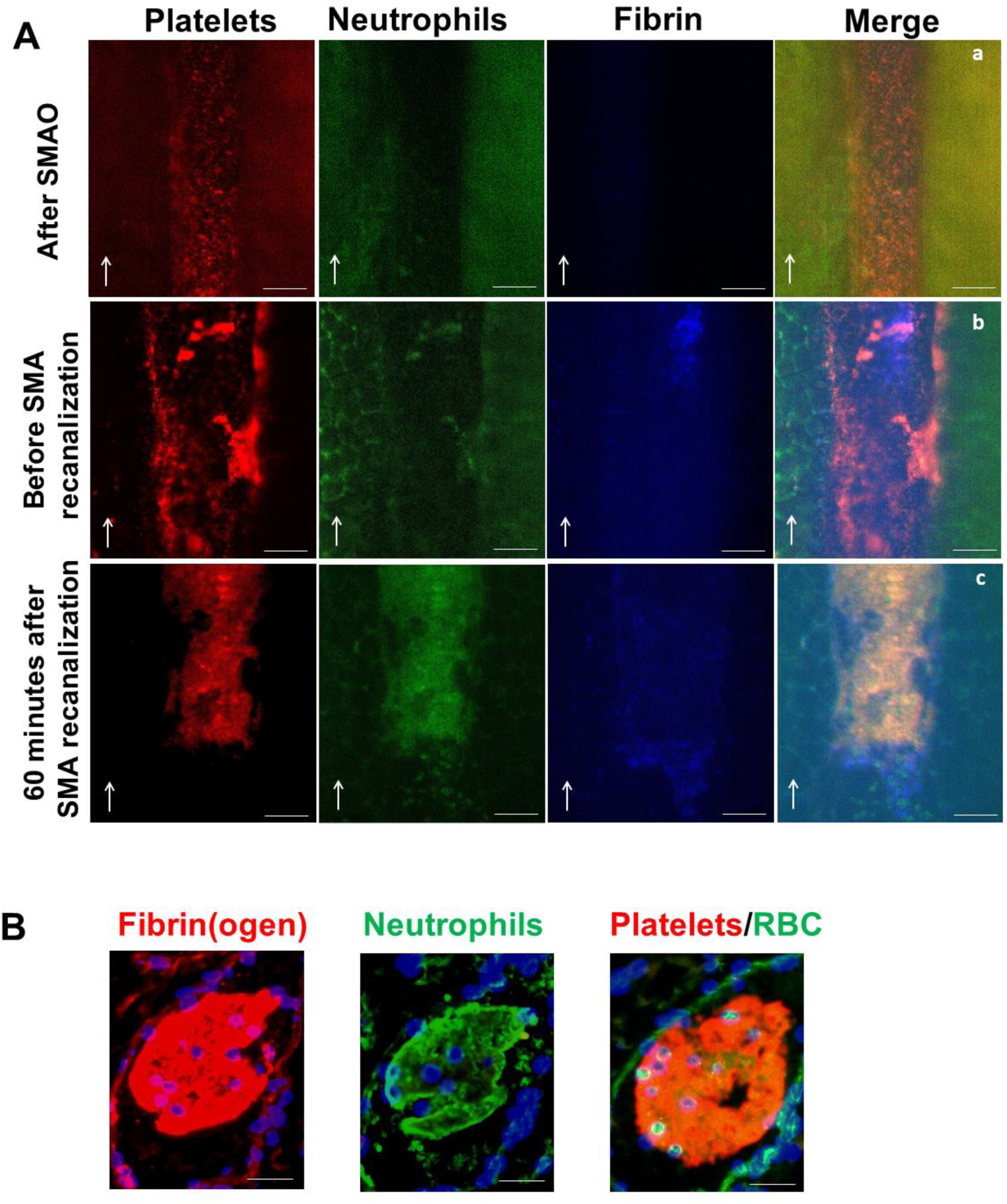
Venular thrombo-inflammatory cell recruitment and thrombus formation. **(A)** Fluorescent intravital microscopy images showing sequential venular recruitment of platelets (anti-GPIX, red), neutrophils (anti-Ly6G, green), and fibrin deposition (anti-fibrin, blue) downstream of the superior mesenteric artery. Images were acquired immediately after SMAO (a), just before arterial recanalization (b), and 60 min after recanalization (c). Arrows indicate blood flow direction. Scale bar, 50 μm. **(B)** Immunofluorescence analysis of venular thrombus sections showing fibrin(ogen) deposition (anti-fibrin(ogen), red), neutrophils (anti-Ly6G, green), platelets (anti-CD42b), and red blood cells. Images are representative of 3–4 independent thrombi. Scale bar, 50 μm.

### Ischemia–reperfusion–induced vascular injury results in intestinal tissue damage

Macroscopic examination of the intestine after mesenteric IR revealed heterogeneous ischemic, hemorrhagic, and necrotic lesions along the small bowel (Figure 6A). Sham mice exhibited preserved macroscopic and microscopic mucosal architecture with intact villi (Figure 6A & 6D). Histological analysis of SMAO intestines showed marked spatial heterogeneity, with preserved areas adjacent to severely damaged regions characterized by epithelial lifting, thinning or loss of the lamina propria, hemorrhage, and leukocyte infiltration (Figure 6D). In SMAO mice, lesions involved 40 ± 3% of the small intestine, with a mean Chiu score of 4, whereas no injury was observed in sham animals (both *p*<0.001; Figure 6B-C). Similar histopathological features were observed in resected human specimens from acute mesenteric infarction, where focal microvascular thrombi were occasionally identified (Figure 7).

**Figure 6:**
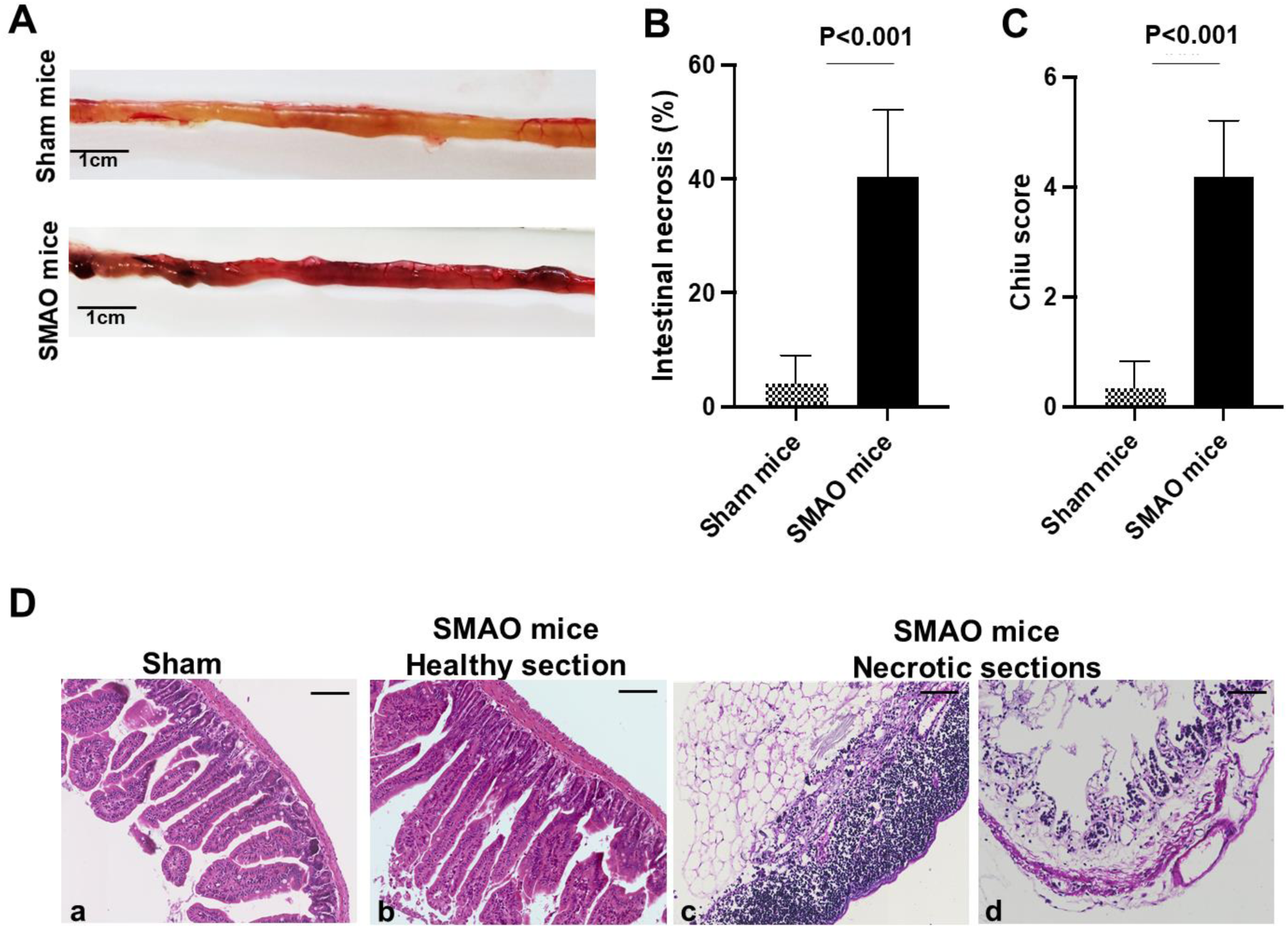
Intestinal injury in experimental and human mesenteric ischemia. **(A)** Representative macroscopic views of the small intestine from sham mice and mice subjected to superior mesenteric artery occlusion (SMAO; 1 h ischemia followed by 1 h reperfusion), illustrating heterogeneous ischemic and reperfusion-associated lesions. **(B)** Quantification of the percentage of necrotic small intestine. **(C)** Histological grading of intestinal injury after ischemia–reperfusion using the Chiu score (0, normal mucosal villi; 5, complete disintegration of the lamina propria with hemorrhage and ulceration). **(D)** Representative hematoxylin–eosin–stained intestinal sections showing preserved mucosa in sham mice (a), adjacent preserved (b) and necrotic areas (c, d) in SMAO mice. Scale bars, 100 μm. Data are presented as mean ± SEM. Images are representative of 3–4 mice per group.

**Figure 7:**
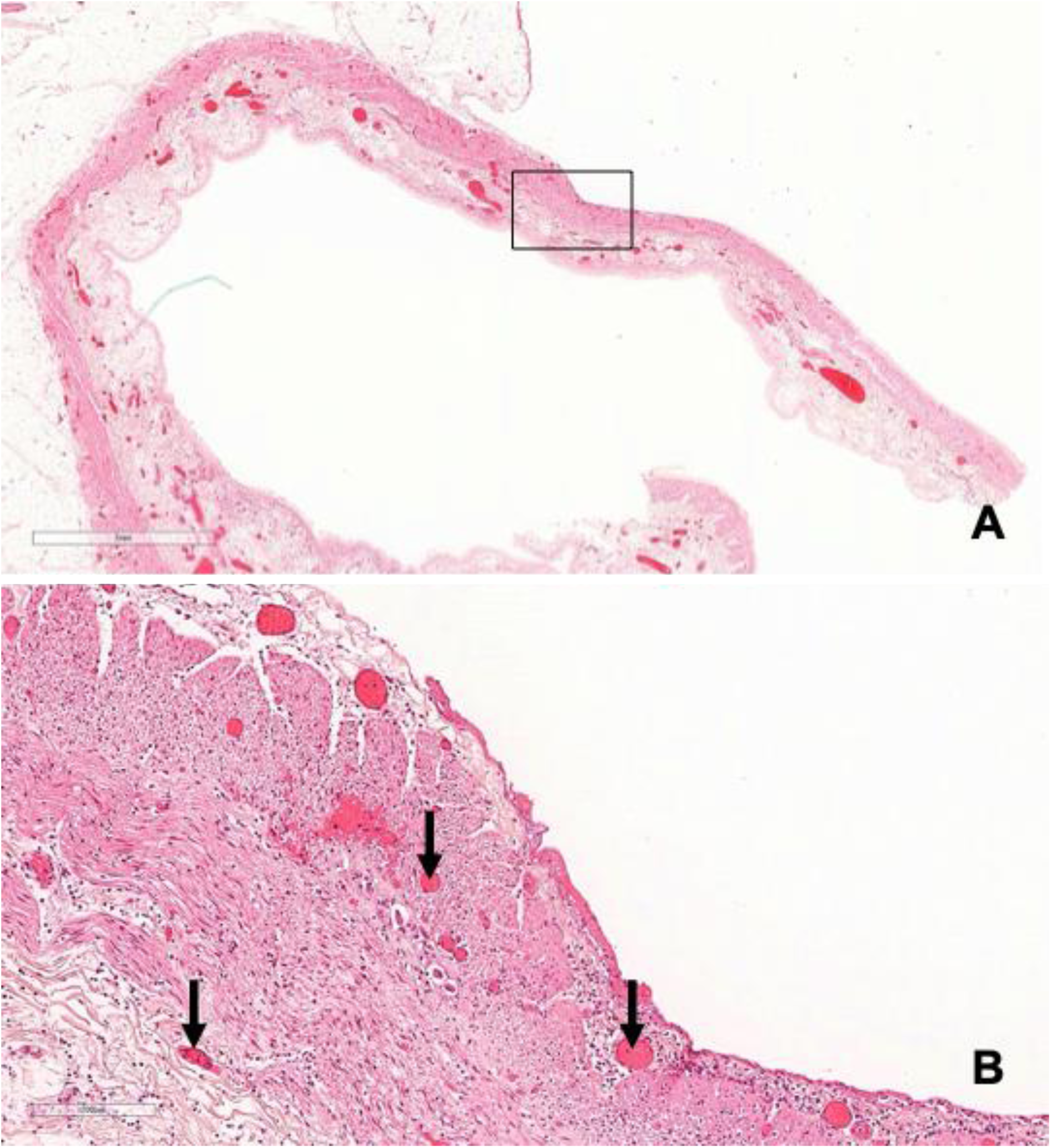
Microvascular thrombosis in human arterial acute mesenteric ischemia. **(A)** Histological section of infarcted small bowel from a patient with acute mesenteric ischemia caused by atherosclerotic occlusion of the proximal superior mesenteric artery, showing extensive transmural ischemic injury involving all layers of the intestinal wall, including the muscularis externa (hematoxylin and eosin staining; original magnification ×10). **(B)** Higher magnification of the boxed area in (A), focusing on the muscularis externa and revealing myocytic necrosis associated with acute microvascular thrombi within the intestinal wall (arrows) (hematoxylin and eosin staining; original magnification ×200).

## DISCUSSION

In this study, we integrated a deeply phenotyped human AMI biobank with a murine superior mesenteric artery occlusion model to bridge clinical observations with mechanistic in vivo analyses. Although arterial acute mesenteric ischemia is characterized by a systemic thrombo-inflammatory response, circulating biomarkers alone do not identify the microvascular origin of these processes. Using intravital microscopy, we revealed a dissociation between arterial and microvascular reperfusion: while arteriolar flow partially recovers after recanalization, venular perfusion remains profoundly impaired, with early blood cell stasis and stable thrombus formation. This venular-centered thrombo-inflammatory response, initiated during ischemia and amplified during reperfusion, provides a mechanistic link between systemic inflammation and persistent microvascular dysfunction in AMI.

Our results are consistent with established thrombo-inflammatory paradigms in which platelets act as early responders to ischemic stress, providing a scaffold for fibrin formation and inflammatory cell recruitment^24^. Neutrophil activation, supported by increased circulating MPO, likely contributes to thrombus maturation and vascular injury^25^. Although neutrophil extracellular traps (NETs) were not directly assessed, their involvement is biologically plausible, as NETs are known to integrate into fibrin networks, stabilize thrombi, and impair thrombolysis in ischemic conditions^26,27^.

The predominance of venular thrombus formation observed in our model is consistent with the low-shear, stasis-prone environment of post-capillary venules, which favors thrombo-inflammatory amplification, whereas higher arteriolar flow limits stable thrombus formation. Importantly, similar venular-predominant microvascular thrombosis despite upstream arterial recanalization has been described in other ischemic organs, particularly in experimental models of acute ischemic stroke. Using intravital microscopy, Desilles and colleagues demonstrated that microvascular thrombosis following arterial recanalization develops primarily within post-capillary venules, despite restoration of proximal arterial flow^28,29^. Thrombi in these models display a platelet-and red blood cell–rich core stabilized by fibrin, with neutrophils preferentially localized at the thrombus periphery. The observation of comparable venular thrombo-inflammatory events in distinct ischemic vascular beds supports the existence of a shared pathophysiological principle governing microvascular failure after large-vessel reperfusion. This convergence strongly supports the translational relevance of our findings and validates the SMAO model.

Microvascular thrombo-inflammation translated into substantial tissue injury. Histopathological analysis demonstrated severe and spatially heterogeneous mucosal damage in SMAO mice, quantified using the Chiu scoring system^23^. Comparable features, including focal microvascular thrombi, were observed in resected human intestinal specimens, supporting the clinical relevance of microvascular involvement in AMI. In humans, however, direct visualization of venular microthrombi is not feasible, and histological analyses are restricted to resected specimens, in which advanced ischemic necrosis often precludes precise characterization of the microvascular compartment involved.

Our results extend previous experimental work showing that restoration of arterial patency does not guarantee sustained microvascular reperfusion. Using laser Doppler flowmetry in a rat ischemia–reperfusion model, Aksoy et al. showed that capillary flow recovers after short ischemic durations but deteriorates following prolonged ischemia despite arterial recanalization^30^. Our intravital microscopy data refine these observations by identifying the venular compartment as a critical site of persistent flow impairment driven by thrombo-inflammatory mechanisms.

Beyond microvascular thrombosis, the intestine displays unique features that may amplify thrombo-inflammation compared with other ischemic organs. Unlike the brain or myocardium, the gut is not a sterile organ and harbors a dense immune cell compartment and complex microbiota. Systemic inflammatory response syndrome (SIRS) is reported in 25–100% of AMI cases^31,32^, whereas ischemic injury involving the heart or brain is associated with SIRS in only 3–22% of cases^33–35^. Intestinal ischemia rapidly induces microbiota alterations, bacterial overgrowth, and translocation of anaerobic species that may escape detection by standard blood cultures (“occult bacteremia”)^36^. Bacterial products and endotoxins further amplify systemic inflammation and may aggravate ischemic injury through impaired epithelial renewal^37^, increased epithelial apoptosis^38^, and microvascular steal phenomena mediated by nitric oxide synthase activation^39^. In line with this concept, experimental models have shown reduced microbial translocation, SIRS, and mortality with oral antibiotics or in germ-free animals^40,41^. Consistently, clinical data support a protective effect of antibiotic strategies in AMI^21,40,41^, motivating the ongoing randomized ORIAMI trial (ORal antibiotics In Acute Mesenteric Ischemia, NCT06387147).

These results may also help reinterpret clinical observations linking vascular comorbidities to worse outcomes. Diabetes has been associated with increased severity of AMI due to mesenteric venous thrombosis^42^, suggesting that pre-existing microvascular disease may amplify venular thrombo-inflammation during ischemia–reperfusion. In arterial AMI, such mechanisms may contribute to the persistently high mortality, which remains in the range of 20–30% even when patients are diagnosed early, managed in expert centers, and treated with prompt arterial revascularization^43^. In this context, the present data provide a biological rationale supporting the current empirical use of anticoagulation in AMI management, while underscoring the limitations of arterial recanalization alone. Importantly, they offer a framework for future pharmacological and interventional studies targeting thrombo-inflammatory pathways, not only in arterial AMI but also across other ischemia–reperfusion settings, including non-occlusive mesenteric ischemia (NOMI) and low-flow states encountered in critical care and major cardiac or vascular surgery. Improved understanding of these downstream processes may also facilitate the identification of novel diagnostic and prognostic biomarkers, another major unmet need in AMI^44^.

In conclusion, mesenteric ischemia–reperfusion induces a venular-centered thrombo-inflammatory response that persists despite arterial recanalization and may contribute to sustained microvascular dysfunction and intestinal injury. By integrating human biomarker profiling with mechanistic intravital imaging, this study provides a unifying framework linking systemic inflammation, microvascular thrombosis, and poor outcomes in AMI, and supports targeting thrombo-inflammation as a therapeutic complement to arterial revascularization.

## Acknowledgments

We thank Olivier Thibaudeau and the HIPNO platform from pathology department of CHU Bichat-Claude Bernard (Paris, France) for assistance with the histology preparation.

## Author Contributions

Study conception and design DF, YB, AN; acquisition of data DF, LK, AB, OC, DCH; statistical analysis DF, AN; drafting of the manuscript: DF, YB, AN; data interpretation and critical revision of the manuscript for valuable intellectual content: DF, VA, MCB, BHTN, YB, AN; study supervision: DF, YB, AN; and patient’s inclusion and care: OC, AN.

## Data sharing

De-identified participant data are available from the corresponding author upon reasonable request, subject to approval by the SURVI/TSUNAMI steering committee and a data-use agreement.

## SUPPLEMENTAL DATA

### METHODS

#### Proteomic serum analysis (untargeted LC–MS/MS)

Serum proteins were first depleted of the 12 most abundant plasma proteins using the Proteome Purify 12 Human kit (R&D Systems) and subsequently concentrated with Amicon Ultra centrifugal filters (Merck Millipore). For each sample, 10 µg of total protein were adjusted to 120 µL of denaturation buffer (4 M urea, 1.5 M thiourea, 50 mM Tris–HCl pH 8.3). Proteins were reduced with 10 mM dithiothreitol for 30 minutes at room temperature and alkylated with 55 mM iodoacetamide for 20 minutes. Samples were digested sequentially, first with 500 ng Lys-C for 3 hours at room temperature, then, after threefold dilution in Milli-Q water, with 500 ng trypsin (Promega) for 16 hours at room temperature. Digestion was terminated by adding formic acid to a final concentration of 3%, and peptide mixtures were stored at –20 °C until analysis.

Peptides were desalted with ZipTip µ-C18 tips (Millipore) before LC–MS/MS analysis on a Q-Exactive Plus mass spectrometer coupled to a Nano-LC Proxeon 1000 system and Easy-Spray ion source (Thermo Scientific). Chromatographic separation was performed using a PepMap C18 precolumn (2 cm, 75 µm i.d., 3 µm) and a Pepmap-RSLC analytical column (50 cm, 75 µm i.d., 2 µm) under a 98-minute gradient from 5% to 35% acetonitrile (0.1% formic acid) at 300 nL/min, for a total run time of 120 minutes. Full MS scans were acquired at 70,000 resolution (m/z 375–1500; AGC 3 × 10⁶), and HCD MS/MS spectra were recorded in Top20 data-dependent mode at 17,500 resolution (AGC 2 × 10⁵; NCE 30%; isolation window 1.4 Da; dynamic exclusion 30 s). Maximum injection times were 50 ms for MS and 45 ms for MS/MS.

Raw data were processed using Proteome Discoverer 2.2 with Mascot 2.5.1. Searches were performed with tolerances of 6 ppm for precursor ions and 0.02 Da for fragments, allowing up to two missed cleavages and considering oxidation (Met), phosphorylation (Ser/Thr/Tyr), carbamidomethylation (Cys), and N-terminal acetylation as variable modifications. Peptide-spectrum matches were validated at a 1% false discovery rate (FDR) using Percolator. Label-free quantification was performed with Progenesis QI 3.0 through chromatogram alignment, normalization on total peptide abundance, statistical analysis of peptide features, and subsequent peptide–protein identification. Protein quantification followed the Hi-3 approach, and only proteins with at least two unique peptides were considered. Differentially abundant proteins were defined by a p-value < 0.05.

